# Evaluate effects of multiple users in collaborative Brain-Computer Interfaces: A SSVEP study

**DOI:** 10.1101/2021.03.05.434173

**Authors:** Tien-Thong Nguyen Do, Thanh Tung Huynh

## Abstract

This study investigates the effects of collaboration on task performance in brain-computer interface (BCI) based on steady-state visually evoked potential (SSVEP). Navigation tasks were performed in a virtual environment under two conditions, e.g., individual performance and team performance. The results showed that average task completion time in the collaborative condition is decreased by 6 percent compared with that of individual performance, which is inline with other studies in collaborative BCI (cBCI) and joint decision-making. Our work is a step forward for the progress in BCI studies that include multi-user interactions.

## I. Introduction

The phenomenon of collective decision-making attracts a lot of interest in the literature recently [1]–[3]. Those researches confirmed that two heads were getting better than one head in the decision-making process, in the case where free communication and interaction are possible between each other. This mechanism is called two-heads-better-than-one (2HBT1) [3]. However, there are still few studies [4]–[6] focusing on the aspect of brain signal decoding. In this study, we investigate this mechanism with a brain-computer interfaces (BCI) system [7]. This work is needed because in a BCI system, typically a game, people tend to play with their teammates. The team performance can be divided into two modes, e.g., collaborative and competitive. In those two modes, users might be encouraged to put more effort, engagement, and attention into the game with their teammates. However, developing such system into the BCI system is still under investigation for the possibility of multiple users in BCI game. Furthermore, the study gives an insight into the impact of multiple users in the cBCI system [8]–[10] compare to individual performances.

One of the first multi-users BCI game reported was Brainball [11]. In this Brainball game, two users have to push a ball one against the other. They are expected to compete by relaxing and their performance is measured by electroencephalography (EEG). The most relaxed player will be the winner. Another study relying on a similar principle can be found in BrainArena [12]. Motor imagery was used to steer a ball into a goal. Eight EEG channels located around the right and left motor cortices were used. This BrainArena game can be played on an individual, collaborative or competitive mode. In another collaborative BCI with event-related potentials (ERP) [6], a large number of participants were asked to participate. Six groups of six people were performing the Go/NoGo task in both offline and online conditions. Their results confirmed that collaboration in BCI could accelerate human decisionmaking. So far, only alpha rhythm [11], motor imagery [12] and ERP [6] have been used for online multi-BCIs. There are still open questions regarding other BCI paradigms, such as SSVEP. In Gurkok’s study [13], they designed an experiment with multi-players BCI based on SSVEP paradigm, but only one participant was playing the game with EEG control. Participant’s partner did not wear an EEG cap, the research target was to check the social impact of collaborative multi-BCI game.

In this study, we further investigated the feasibility of multiple users in SSVEP based BCI in cBCI mode in Virtual Environment game. Based on experience with collaborative BCI for accelerating human decision-making [14] and other studies with motor imagery and alpha rhythm [11], [12], we sought to imitate the design using a SSVEP paradigm.

## II. Method

### A. Participants

Ten graduate students (5 females: mean age 25.3±0.8 years old ranging from 24 and 26) voluntarily participated in this experiment. They all had normal or corrected to normal vision. No psychiatric disorders were reported from themselves and their family members. Members of each pair knew each other.

### B. Experimental Design

*SSVEP technique* SSVEP is a synchronized brain response induced by a repetitive visual stimulus, flickering at a constant frequency [15]. The principle is same for different kinds of stimuli types, oscillating or moving image with constant frequency (stimulus frequency) will elicit brain response to the same stimulus frequency and its harmonics.

There are several stimuli types with different level of complexity ranging from simple to complex (figure 1) [15]

- goggles [16]
- light-emitting diodes (LED) [17]
- monitor screen: liquid crystal display (LCD) or cathode ray tube (CRT)
  – simple: just black-white color
  – complex pattern: checkerboard [18], emotional pictures [19]
  – steady-state motion visual evoked potentials [20]
  – moving grating
  – gaussian field [21]
  – intensity of the monitor luminance [22]

**Fig. 1.**
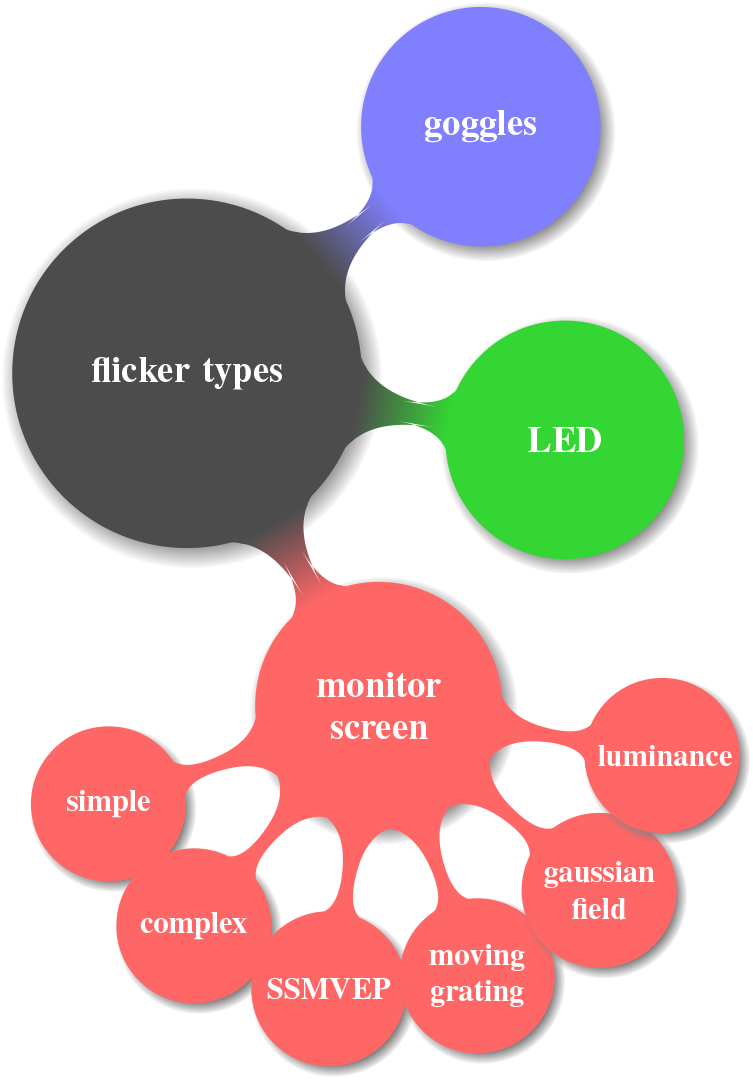
Flicker types for SSVEP

In this study, we chose the intensity of the monitor luminance to generate flicker. This method can be adapted and incorporated into the monitor, which can be more flexible for user feedback. The user interface was created with OpenGL embedded in Psychophysics Toolbox-3 for MATLAB [23] (figure 2). Through this user interface, the participant can control the ball’s movement with three commands: forward, left, and right. This game’s mission is to move forward the ball while avoiding obstacles and reach the destination as fast as possible. The completion time was measured as user performance.

**Fig. 2.**
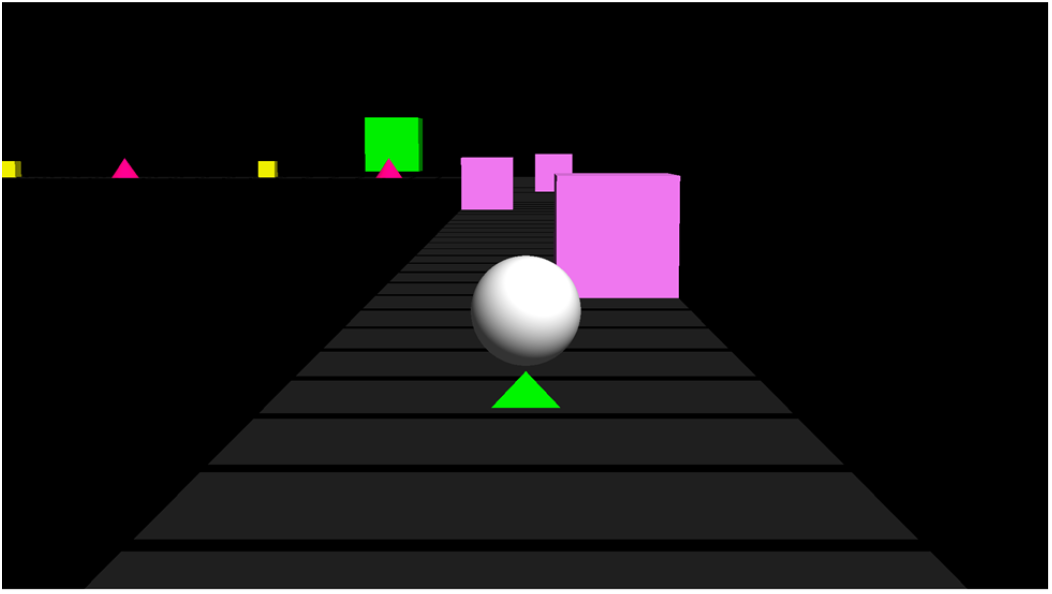
The experimental user interfaces.

### C. Experimental procedure

Figure 3 describes the procedure of our experiment. In the collaborative BCI task, five dyads of participants (two participants x five teams) wore EEG caps, and their data were recorded simultaneously. Participants sat in a comfortable chair with armrests and were given instructions regarding the experimental procedure. A 32 multi-channel EEG amplifier (BrainAmp by Brain Products, Munich, Germany) was used to record the EEG signal. The data were recorded at 1000 Hz with a band-pass filter of 0.5-50 Hz. Eight EEG channels were recorded in the occipital area: PO3, PO4, PO7, PO8, POz, O1, O2, and Oz, according to the international 10-20 system (Figure 4). The nose and AFz were used as reference and ground, respectively. The impedance was kept below 10 kΩ throughout the experiment.

**Fig. 3.**
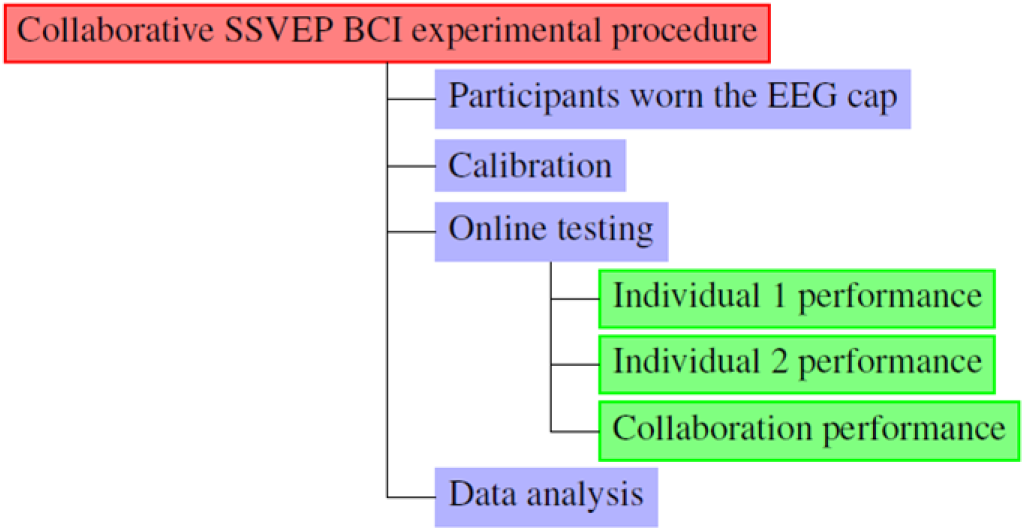
Experimental procedure.

**Fig. 4.**
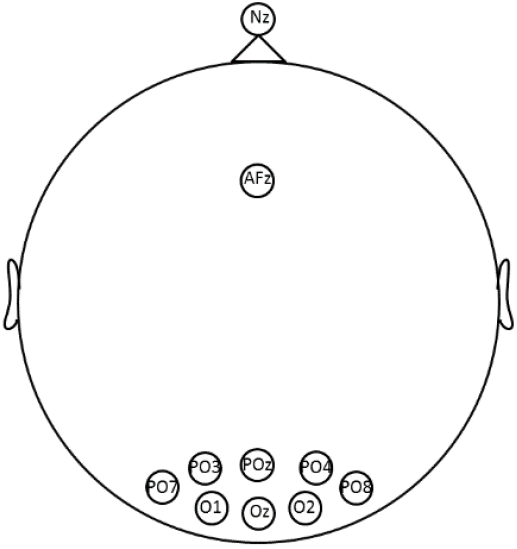
The EEG electrode channels were used in this experiment.

Participants were instructed to pay attention to a square filled with a high arousal picture in the calibration session. The emotional picture continuously oscillated at a frequency of 6 Hz, 7.5 Hz, and 10 Hz, respectively. Twenty trials were recorded for each case (20 trials for the 6 Hz, 20 trials for the 7.5 Hz, and 20 trials for the 10 Hz condition), and each trial lasted for 12 s. To minimize carryover effects, an interval of 4s was used between trials, and a resting time of 10 s was allocated between each frequency condition. It took about 15 minutes for each participant to complete the calibration session. Following the calibration session, optimal parameters were chosen by the canonical correlation analysis (CCA) algorithm.

Figure 5 shows the schematic design for this experiment. Generally, the frameworks of both schemes are same and have three main steps:

- Brain signal acquisition from Brain vision hardware.
- CCA algorithm for classifying user intention. The output of the classifier was used as the input for virtual environment game. Both virtual environment and classifier communicate with each other by user datagram protocol (UDP).
- Virtual environment plays the role of visual feedback.

**Fig. 5.**
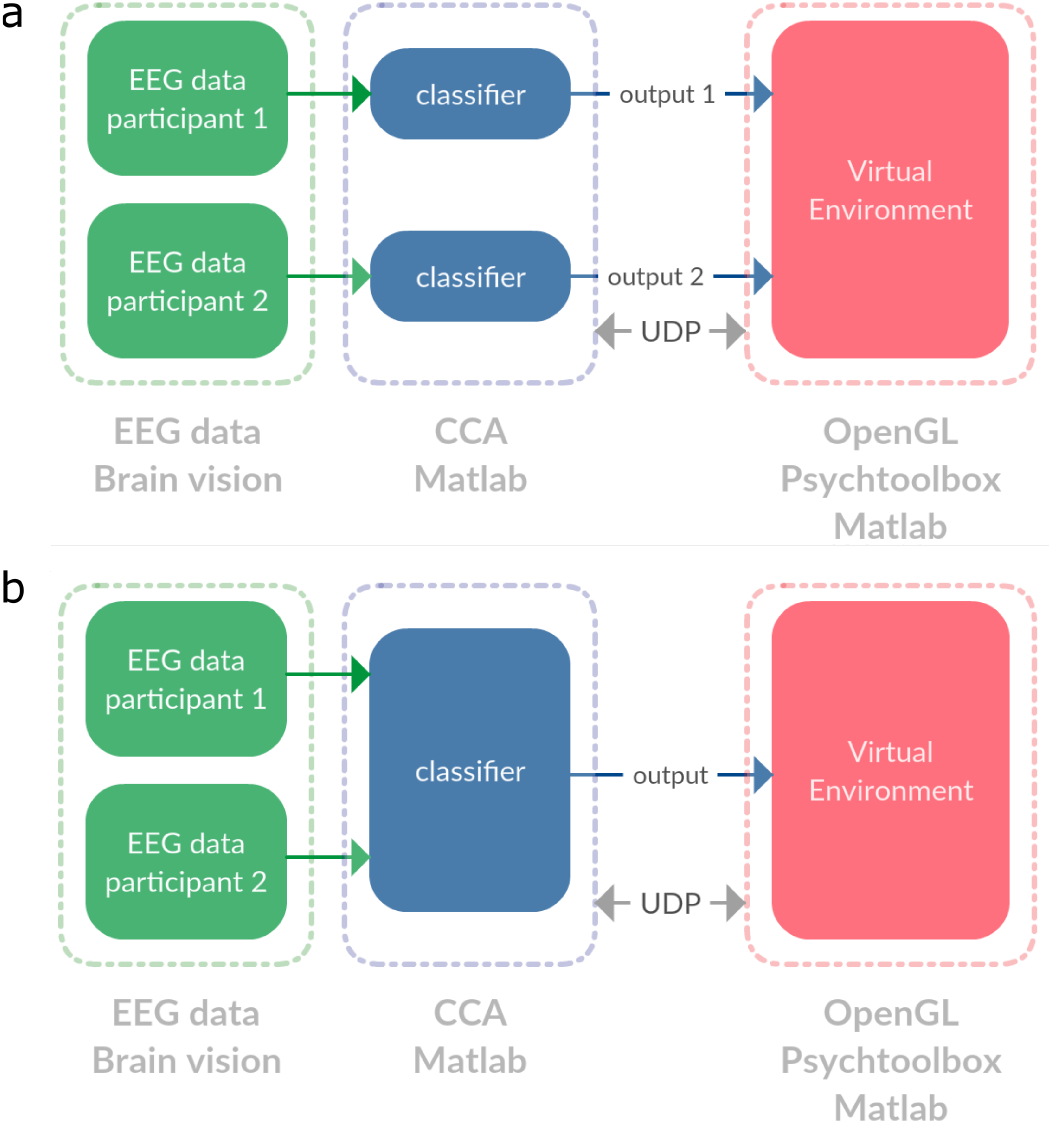
Collaborative scheme setup for (a) type 1 and (b) type 2, respectively.

In addition, two different strategies are considered for analyzing SSVEP signal of multiple players:

- Multiple input commands for Virtual Environment (figure 5a). The EEG data of each participant was classified separately to give out two independent commands to control the ball movement in the virtual environment.
- Single input command for the Virtual Environment (figure 5b). Both EEG user data from Brain vision hardware were merged together and classified by CCA to give out the single input for virtual environment.

### D. Data processing

In this study, we used CCA method for the classification [24], [25]. CCA is a machine learning method that enables to find an underlying correlation between two sets of variables [26]. For two sets of variables X and Y, CCA method consists of finding vectors w and v that maximize the correlation between *w^T^*X and *v^T^*Y.

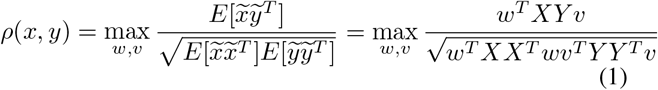

where *ρ* indicates the maximum correlation coefficient, X refers to the set of multi-channel EEG data signals, and Y is reference signals set that have the same length as X:

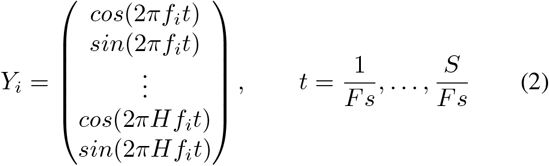

H is the number of harmonics, the classifier’s output is recognized as:

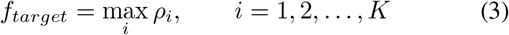

## III. Results

### A. Behavioral performance

As shown in the figure 6, figure 7, participants showed better performance in the collaboration condition compared to those in the individual conditions. The collaboration results tended to show a reduced completion time of up to 6 percent. While earlier studies have reported increased performances for collaboration-based multiuser BCIs in ERP-based [6], or MI-based paradigms [12], here we demonstrate the evidence of a positive effect of synchronizing collaboration for online SSVEP-based BCI.

**Fig. 6.**
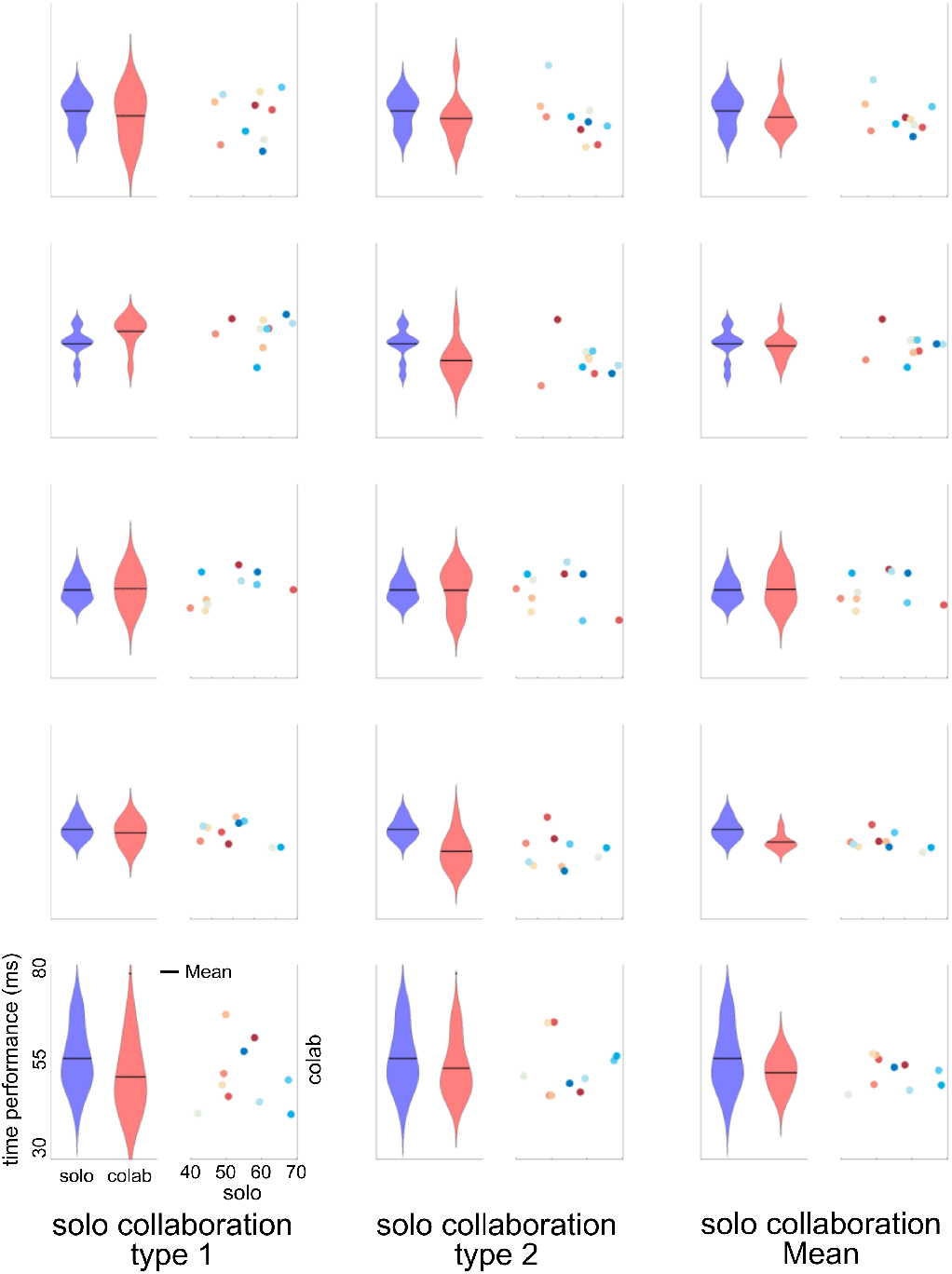
Time performance for each pair.

**Fig. 7.**
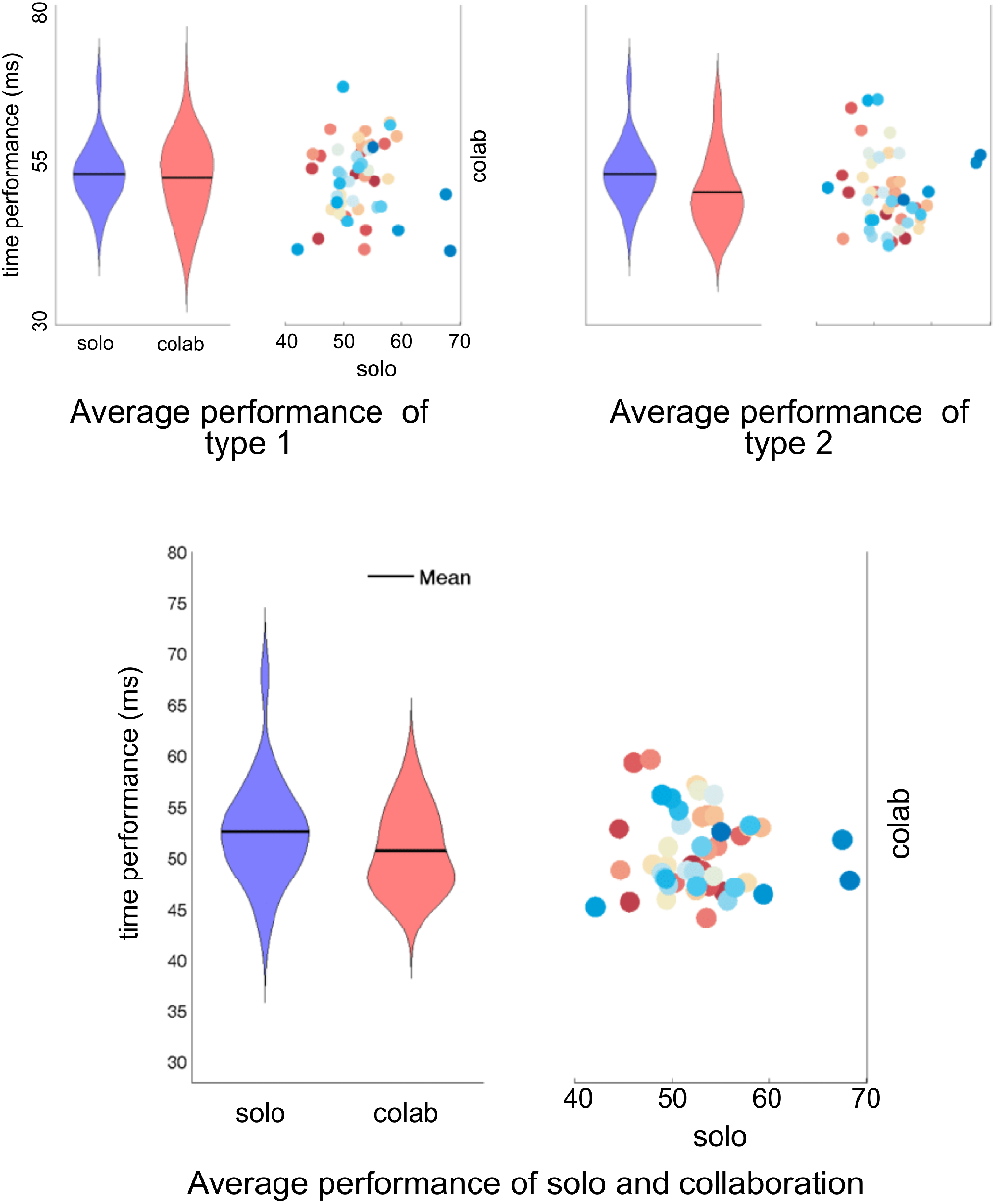
The average of time performance.

### B. EEG data analysis

The more channels we used for classifying data under the machine learning aspect, the more accuracy we might get. This trend can be seen clearly in the figure 8, and figure 9. On the other hand, under the enjoyment aspect illustrated in figure 7, we could see that when people performed collaborative task, they tend to be more involved than individual performance.

**Fig. 8.**
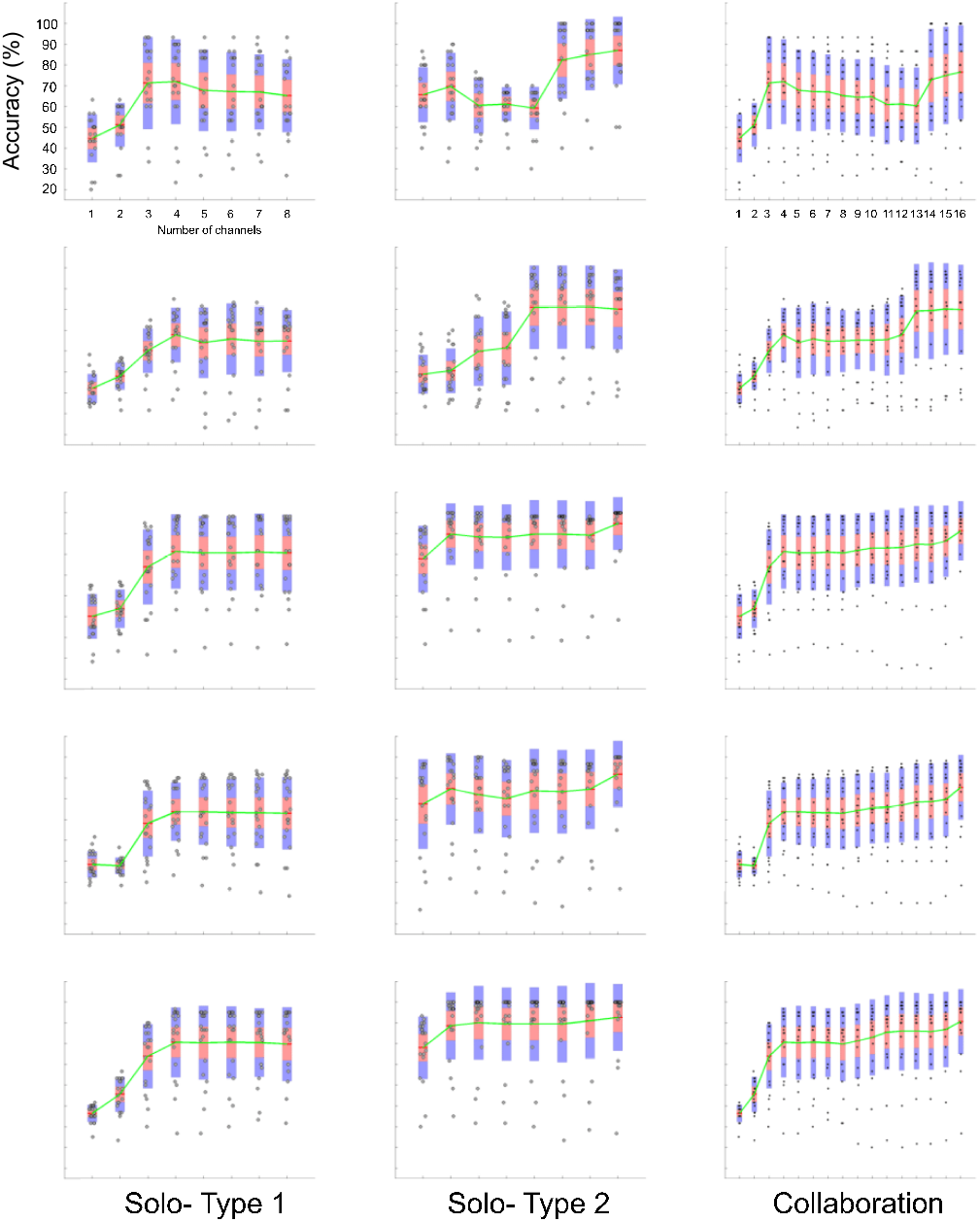
Classifier accuracy of each pair.

**Fig. 9.**
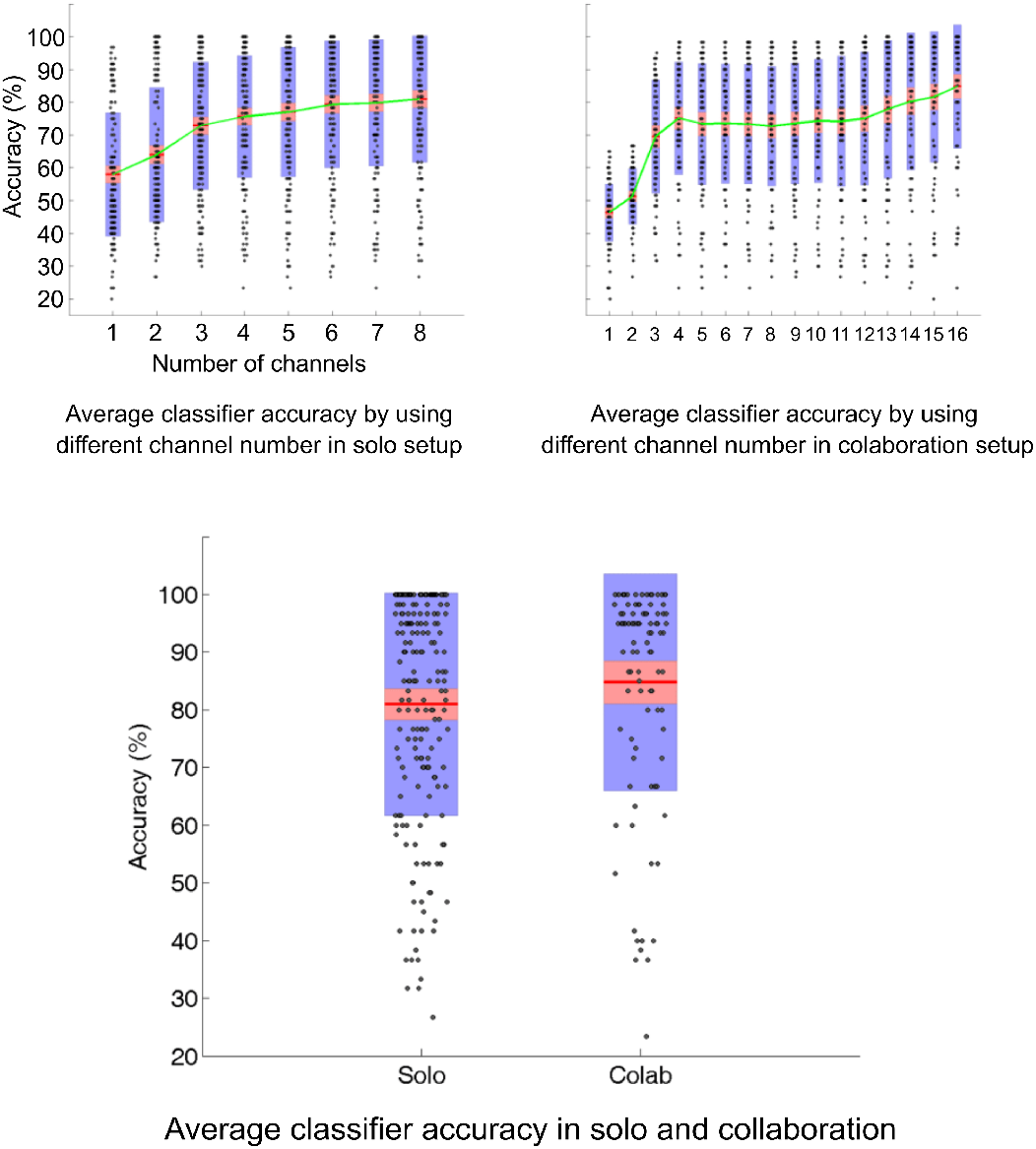
The average of classifier accuracy.

## IV. Discussion and Conclusions

Our work showed that multiple users could benefit from a synergetic effect of collaboration compared with individual performance in terms of brain signal decoding. It also provides another solution for end-users interaction with the BCI system. However, the present work lacks a comparison of a larger majority of machine learning techniques for classifying user intention that should be considered for future works. Another factor that needs to be considered is system response time, as it is a trade-off with the classifier accuracy. Finally, the adaptive user-dependant factor could be examined based on the confidence levels and flicker frequency. For instance, a participant with higher confidence can have a higher weight factor in the classifier.

